# c-Jun Signaling During Initial HSV-1 Infection Modulates Latency to Enhance Later Reactivation in addition to Directly Promoting the Progression to Full Reactivation

**DOI:** 10.1101/2023.11.10.566462

**Authors:** Sara A. Dochnal, Abigail L. Whitford, Alison K. Francois, Patryk A. Krakowiak, Sean Cuddy, Anna R. Cliffe

## Abstract

Herpes simplex virus-1 (HSV-1) establishes a latent infection in peripheral neurons and can periodically reactivate to permit transmission and clinical manifestations. Viral transactivators required for lytic infection are largely absent during latent infection and therefore HSV-1 relies on the co-option of neuronal host signaling pathways to initiate its gene expression. Activation of the neuronal c-Jun N-terminal kinase (JNK) cell stress pathway is central to initiating biphasic reactivation in response to multiple stimuli. However, how host factors work with JNK to stimulate the initial wave of gene expression (known as Phase I) or the progression to full, Phase II reactivation remains unclear. Here, we found that c-Jun, the primary target downstream of neuronal JNK cell stress signaling, functions during reactivation but not during the JNK-mediated initiation of Phase I gene expression. Instead, c-Jun was required for the transition from Phase I to full HSV-1 reactivation and was detected in viral replication compartments of reactivating neurons. Interestingly, we also identified a role for both c-Jun and enhanced neuronal stress during initial neuronal infection in promoting a more reactivation-competent form of HSV-1 latency. Therefore, c-Jun functions at multiple stages during HSV latent infection of neurons to promote reactivation. Importantly, by demonstrating that initial infection conditions can contribute to later reactivation abilities, this study highlights the potential for latently infected neurons to maintain a molecular scar of previous exposure to neuronal stressors.

**Importance:** The molecular mechanisms that regulate the reactivation of HSV-1 from latent infection are unknown. In addition, studies on the mechanisms of reactivation can be complicated by factors that act during latency establishment to ultimately impact reactivation. Here we focused on the host transcription and pioneer factor c-Jun because it is the main target of the JNK cell stress pathway known to be important in exit of HSV from latency. Surprisingly, we found that c-Jun does not act with JNK during exit from latency but instead promotes the transition to full reactivation. Moreover, c-Jun and enhanced neuronal stress during initial neuronal infection promoted a more reactivation-competent form of HSV-1 latency. c-Jun therefore functions at multiple stages during HSV latent infection of neurons to promote reactivation and serves as a relevant therapeutic target for attenuating reactivation and related clinical consequences. Importantly, this study contributes to a growing body of evidence that *de novo* HSV-1 infection conditions can modulate latent infection and impact future reactivation events, raising important questions on the clinical impact of stress during initial HSV-1 acquisition on future reactivation events and consequences.

## Introduction

Long-term viral infection can be regulated at multiple steps by the activation of host-cell signaling pathways. The Herpesviruses establish lifelong latent infections within specific host cells and periodically reactivate to permit transmission. During latent infection, the double-stranded DNA herpesvirus genomes reside in an epigenetically silent state, coated with host histone proteins within host nuclei (1–6). The transcription of viral lytic mRNAs is largely repressed and therefore viral proteins that mediate lytic gene transactivation during productive infection are largely absent. To reactivate, repressed herpesviruses rely on host signaling pathways that appear to play important physiological roles in their respective latent cell type. For example, the Gammaherpesvirus Epstein-Barr virus (EBV) establishes latency in memory B cells, and reactivation can be initiated following B cell receptor ligation and activation (7). Likewise, the Betaherpesvirus human cytomegalovirus (HCMV) establishes a latent infection in hematopoietic progenitor cells and reactivation can be initiated through cytokine signaling, including TNFɑ, which mediates differentiation (8–11). Following activation of these signaling pathways, host transcriptional factors activated downstream of signaling pathways can bind viral genomes to initiate lytic expression (12). Understanding the contribution of host cell proteins to the initiation of reactivation can help uncover the mechanisms of viral expression and ultimately identify targets for therapeutics that can prevent the occurrence of reactivation.

The Alphaherpesvirus, Herpes Simplex Virus-1 (HSV-1), establishes a latent infection in post-mitotic neurons, during which lytic promoters are assembled into heterochromatin, as defined by the enrichment of trimethylated histone H3 lysine 27 (H3K27me3) and di- and tri-methylated lysine 9 (H3K9me2/3) (13–18). Periodically, following common physiological stressors like fever, UV exposure, and psychological stress, full reactivation, as characterized by new infectious virus production, can occur. At the neuronal level, the activation of certain signaling pathways have been found to induce reactivation (reviewed in (19)). These include the loss of neurotrophic factor support (20–27), enhanced neuronal hyperexcitability (21, 28), and perturbation of the DNA damage response and axonal damage (29, 30). However, unlike the studies of EBV and HCMV, the host transcription factors that act downstream of a reactivation stimulus are largely unknown.

Transcriptionally, full reactivation of HSV-1 mirrors *de novo* acute replication, which takes place in neuronal and non-neuronal cells; viral immediate early (IE) gene transcription precedes and is essential to early (E) gene expression, which is required for viral DNA (vDNA) replication, and subsequent late (L) gene transcription. Efficient IE gene expression and therefore the entire downstream lytic transcriptional cascade during acute infection or full reactivation, requires viral trans-activator VP16 (31–34). VP16 complexes with cellular factors involved in transcriptional activation, including general transcription factors, ATP-dependent chromatin remodelers, RNA polymerase, and histone-modifying enzymes that may remove repressive H3K9me2/3 and add euchromatin-associated modifications (35–40). In the context of acute infection, VP16 is delivered to the host cell nuclei with the incoming virus tegument. However, during a latent infection, transcription of the gene encoding VP16 (*UL48*) is restricted and therefore viral gene expression must initiate in the absence of VP16 protein by alternative host or viral factors.

There is accumulating evidence that reactivation is a biphasic process. Phase I gene expression precedes “full reactivation” (also referred to as “Phase II”) as a transcriptional burst of all classes of lytic viral genes, with late gene expression being uncoupled from viral DNA replication. This Phase I gene expression phenomenon has been observed in both *in vitro* and *ex vivo* models of HSV reactivation (21, 25, 28, 31, 41, 42). The use of *in vitro* model systems has enabled the molecular mechanisms of Phase I and Phase II reactivation to be further teased apart. In these models, Phase I reactivation does not require VP16 nor activation of the host histone demethylase enzymes that remove H3K27me3 and H3K9me2 (21, 25, 28, 31, 41). However, Phase I reactivation, and as well as the progression to full Phase II, requires the activation of cellular c-Jun N-terminal kinase (JNK), which is specifically redirected to perform a neuronal stress response by mixed lineage kinase protein dual leucine kinase (DLK) and the JNK scaffold protein, JNK-interacting protein-3 (JIP3). The contribution of this neuronal stress signaling pathway was first demonstrated during reactivation using small molecule inhibitor LY294002, which inhibits the activation of the PI3-kinase and AKT-pathways that occur downstream of NGF-signaling (25). Since this discovery, the requirement of DLK/JNK for viral reactivation has been demonstrated using multiple systems and triggers converging upon diverse cellular pathways. HSV-1 can co-opt an innate immune signaling pathway mediated by IL-1, which induces neuronal hyperexcitation and subsequent reactivation (28). HSV-1 reactivation is also elicited via disruption of neuronal DNA damage or repair pathways, for example through the addition of DNA damaging agents or the inhibition of the ATM-dependent repair pathway by AKT inhibition (43, 44). In response to a combinatorial stimulus of an NGF-deprivation mimic (LY294002), neuronal hyperexcitability (forskolin), and heat shock, reactivation can be induced from a very silent form of latency established *in vitro* (21). DLK is integral to reactivation using all these stimuli, and JNK has also been demonstrated to be required when tested in these systems.

JNK activation during Phase I of HSV-1 reactivation results in the phosphorylation of serine (S10) neighboring repressive mark H3K9me3, and possibly H3K27me3, on the viral genome during Phase I (25). This phospho/methyl switch is known to result in the eviction of repressive reader proteins and therefore likely permits a chromatin environment that is conducive to transcription (45–47). However, additional host proteins, including transcription and pioneer factors, that directly promote viral gene expression would also be required for Phase I reactivation. Moreover, JNK lacks DNA binding capabilities, which suggests that an additional DNA-binding protein mediates JNK recruitment to viral chromatin.

DLK-mediated activation of JNK both up-regulates and phosphorylates its primary transcriptional factor target c-Jun, which can mediate neuronal cell death and axon pruning following the loss of nerve growth factor signaling (48–53). We previously observed c-Jun activation in neuronal models used for HSV-1 latency following PI3-kinase inhibition (25) and forskolin treatment (28). Unlike traditional transcription factors, c-Jun can maneuver or pioneer through heterochromatin to modulate genome accessibility in a broad range of host cells, including neurons (54, 55). We therefore set out to investigate whether c-Jun, is critical for HSV-1 reactivation, with the hypothesis that c-Jun up-regulation and phosphorylation by JNK acts to induce Phase I gene expression. Interestingly, we found that c-Jun protein was required for reactivation during Phase II, but does not function directly during Phase I. In carrying out this study we also found that activation of c-Jun via neuronal stress during *de novo* infection resulted in enhanced reactivation. Therefore, we show that cell stress, and potentially other signals that could activate c-Jun during initial neuronal infection, can have a long-term impact on either the neuron or viral chromatin to regulate the propensity of HSV-1 to reactivate.

## Results

### c-Jun depletion prior to HSV-1 latency establishment restricts both Phase I gene expression and full reactivation

To Investigate the contribution of c-Jun to HSV-1 latency and reactivation, we used an *in vitro* primary neuronal model because this permits the easy manipulation of c-Jun at different times during the latency/reactivation cycle. In addition, robust reactivation can be achieved in intact neurons using this system. Latent infection was established in sympathetic neurons isolated from the superior cervical ganglia (SCG) of newborn mice as described previously (21, 25, 28, 56). Neurons were infected with a gH-null virus containing Us11-GFP (Stayput-GFP), which permits the quantification of individual reactivating neurons neuronal model (21). Depletion of c-Jun protein was validated in SCG neurons using three independent shRNA lentiviruses (Fig 1A). c-Jun was depleted from primary neurons using the two most effective lentiviruses (sh-cJun2 and sh-cJun3) and subsequently infected with HSV-1 Stayput-GFP at an MOI of 7.5 PFU/cell in the presence of acyclovir (ACV; 50μM) for six days. Two days post removal of the acyclovir, the infected neurons were reactivated with the PI3-kinase inhibitor, LY294002 (20μM), and the number of Us11-GFP-positive neurons was quantified at 48 hours post-treatment, which is indicative of Phase II reactivation, in addition to Phase I lytic gene expression analysis at 18 hours post-treatment. Despite similar viral DNA loads during latency (Fig 1B), both full reactivation (Fig 1C, F) and Phase I gene expression (Fig 1D-E, G-H) were significantly reduced in c-Jun depleted cultures. Therefore, the presence of c-Jun protein during HSV latent infection was required for the initial exit from latency and Phase I gene expression.

**Figure 1:**
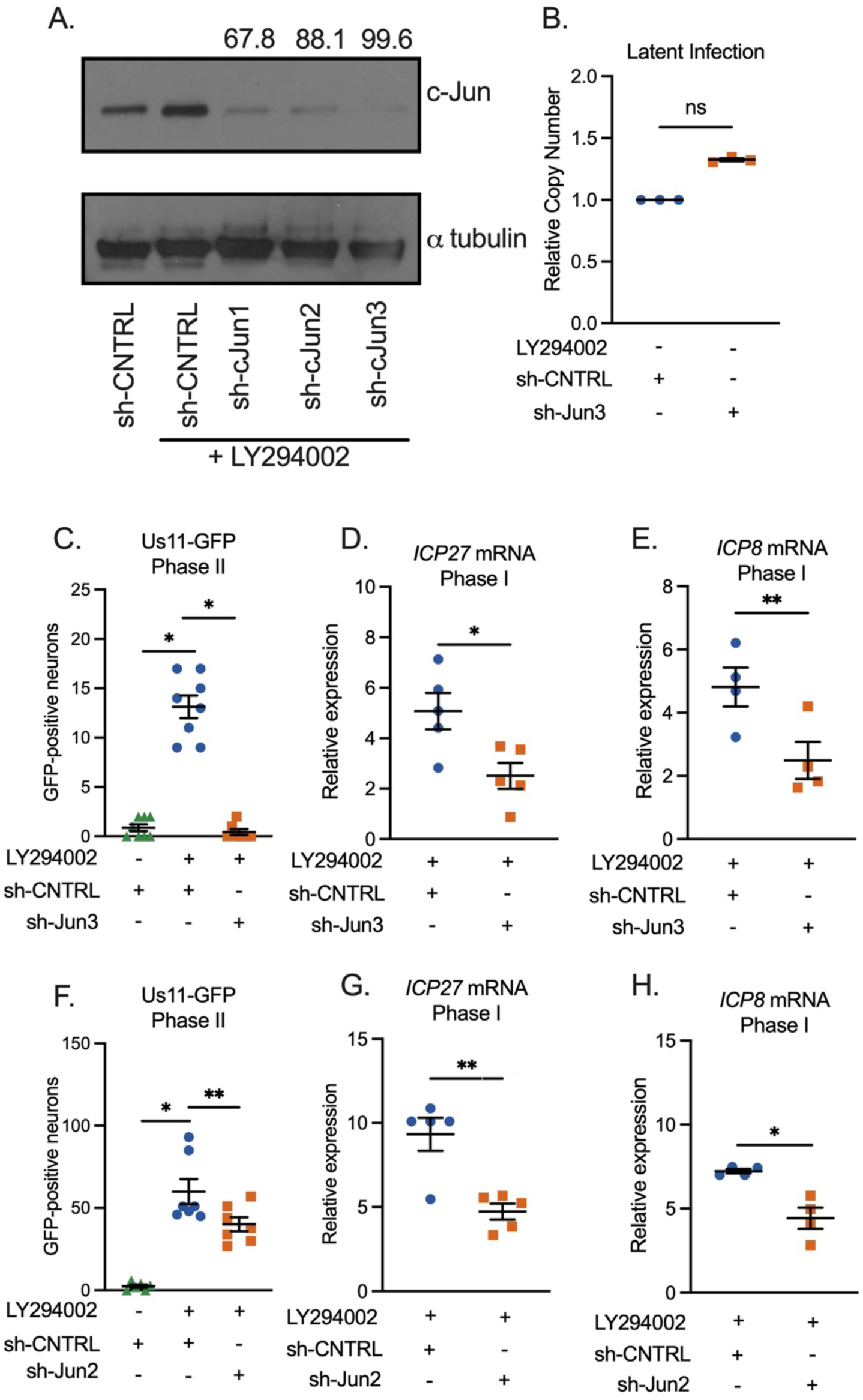
c-Jun depletion prior to latency establishment limits both Phase I gene expression and full reactivation. A) Neurons were transduced with a non-targeting shRNA control lentivirus or one of three independent lentiviruses expressing shRNAs that target Jun (sh-cJun1, sh-cJun2, sh-cJun3). 5 days post transduction, LY294002 was added to neurons for 18 hours and western blotting for c-Jun or α-tubulin was performed. The percentage knockdown of c-Jun normalized to α-tubulin is shown. B-H) Following c-Jun depletion with sh-cJun3 (B-E) or sh-cJun2 (F-H), primary neurons were infected with Stayput-GFP at an MOI of 7.5 PFU/cell in the presence of acyclovir (ACV; 50 μM) for six days and then reactivated two days after the removal of acyclovir with LY294002 (20 μM). B) Quantification of relative latent viral DNA load at 8 days post-infection. Biological replicates from 3 separate dissections C & F) Quantification of the number of GFP-positive neurons at 48 hours post-stimulus. Individual biological replicates from at least 3 individual dissections. D-E, G-H) Relative viral gene expression at 18 hours post-stimulus compared to latent samples quantified by RT-qPCR for ICP27 (D,G), ICP8 (E,H) normalized to cellular control mGAPDH. Statistical comparisons were made using normal or non-normal (Wilcoxon, B, C, F) Paired t-Test. Biological replicates from at least 3 individual dissections. Individual biological replicates along with the means and SEMs are represented. * P < 0.05; ** P < 0.01.

### c-Jun depletion following latency establishment does not prevent entry into Phase I

The data shown in Figure 1 support our hypothesis that c-Jun mediates Phase I gene expression and ultimately full reactivation downstream of the DLK/JNK signaling cascade. However, these experiments come with the caveat that c-Jun was depleted prior to HSV-1 infection and therefore we cannot rule out an indirect role of c-Jun in latency establishment that ultimately perturbs reactivation. To test the role of c-Jun solely in Phase I reactivation, the time point where both DLK and JNK act to promote exit from latency, and therefore reactivation, we depleted c-Jun following HSV-1 infection and latency establishment and analyzed Phase I gene expression. In contrast to what we observed following depletion of c-Jun prior to infection, Phase I gene expression was not perturbed following c-Jun depletion after latency establishment (Fig 2A-C). Importantly, we verified knockdown at the RNA level for each replicate (Fig 2D). This phenotype was not limited to a single shRNA clone as Phase I gene expression remained unchanged with the additional two independent c-Jun lentiviruses (Fig 2E-G) that were previously validated (Fig. 2A). In addition, this phenotype was not limited to one trigger as c-Jun was also not required for Phase I gene expression triggered by forskolin (Fig 2H-K). In contrast, c-Jun depleted cultures displayed unexpectedly enhanced Phase I gene expression following treatment with forskolin. Therefore, our data indicate that c-Jun does not play a direct role alongside DLK and JNK in supporting Phase I gene expression during reactivation. In addition, these data contrasted what was observed for the impact of c-Jun depletion pre-infection on entry into Phase I gene expression.

**Figure 2:**
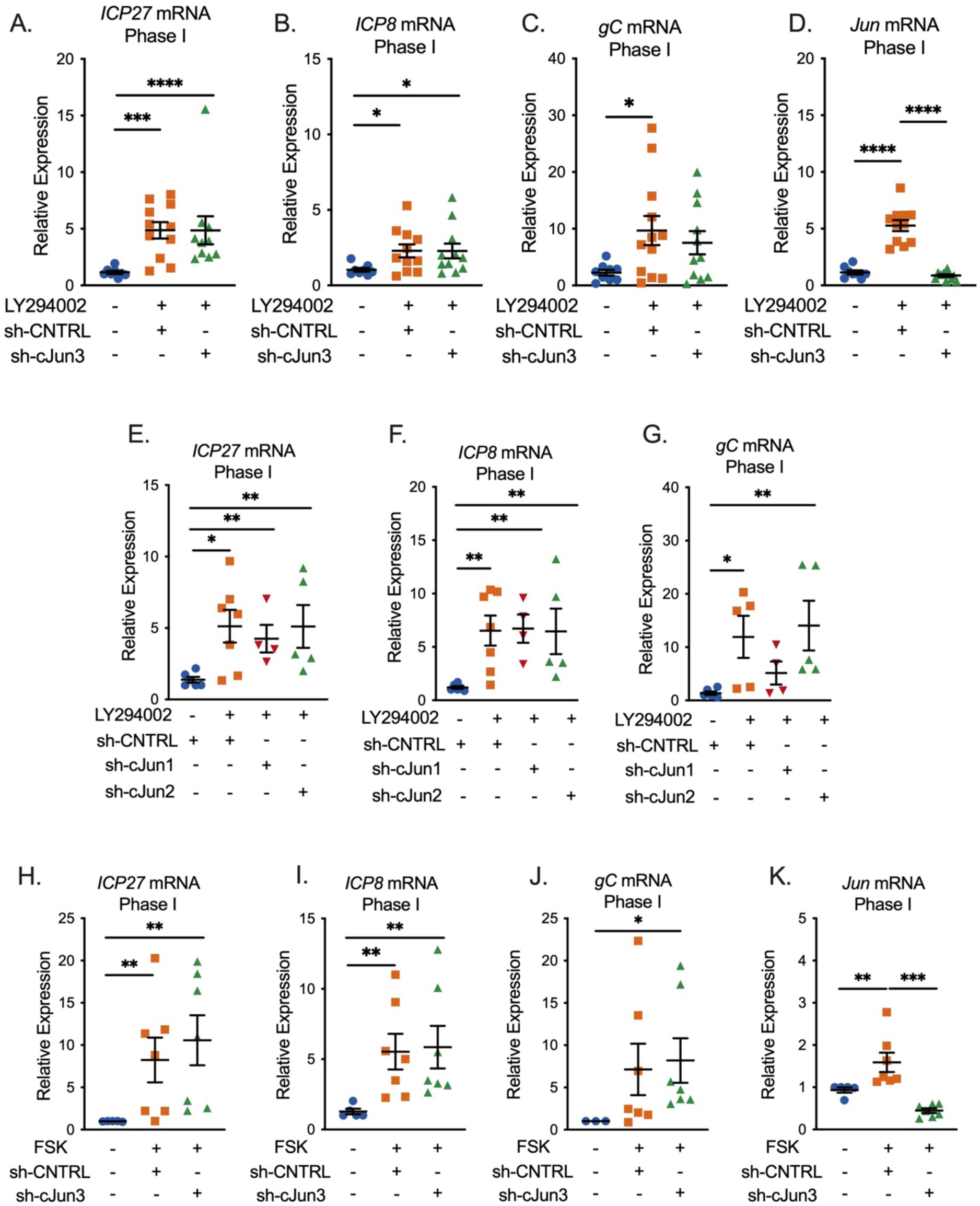
c-Jun is not necessary for Phase I gene expression. (A-K) Latently infected neurons were transduced with a non-targeting shRNA lentivirus or sh-cJun3, sh-cJun2, or sh-cJun1 at 6 days post-infection and reactivated 5 days later. In (A-G) neurons were reactivated with LY294002 and in (H-K) with forskolin. RT-qPCR was carried out at 18 hours post-reactivation for; ICP27 (A, E, & H), ICP8 (B, F, & I), gC (C, G, & J) and cellular Jun (D&K) at 18 hours post-stimulus is represented. N=6 biological replicates from at least 3 independent dissections. Statistical comparisons were made using non-normal (Mann-Whitney) t-Test. Individual biological replicates along with the means and SEMs are represented. * P < 0.05; ** P < 0.01.

### c-Jun is Required for the Expression of Late Genes during Full (Phase II) Reactivation

Although we did not detect a role for c-Jun in entry into Phase I gene expression, we went on to examine whether it played any role during HSV-1 reactivation. We therefore again depleted c-Jun prior to reactivation and quantified entry into full, Phase II reactivation. We now did detect a role for c-Jun during Phase II/full reactivation, as the numbers of Us11-GFP positive neurons were significantly decreased following c-Jun depletion (Fig 3A). This was verified using a second c-Jun shRNA lentiviral clone (Fig 3G). This indicated that c-Jun was required for Us11 expression. Because Us11 is a true late gene, we additionally quantified immediate early (Fig 3B), early (Fig 3C), and late viral transcripts (Fig 3D-E) during Phase II. Importantly, c-Jun was specifically required for expression of the late genes tested and not the immediate early gene ICP27 nor the early gene ICP8. The impact of c-Jun depletion on late gene expression was validated using a second c-Jun shRNA clone (Fig 3H &I). We also quantified the impact of c-Jun depletion on the progression to full reactivation induced by forskolin. Consistent with the data for PI3-kinase induced reactivation, the numbers of Us11-GFP positive neurons were decreased in the c-Jun depleted neurons, and this was verified using two independent shRNA clones (Fig 3F,J). Therefore, these data indicate that c-Jun is directly required for full, Phase II HSV reactivation and acts specifically on late viral transcripts.

**Figure 3:**
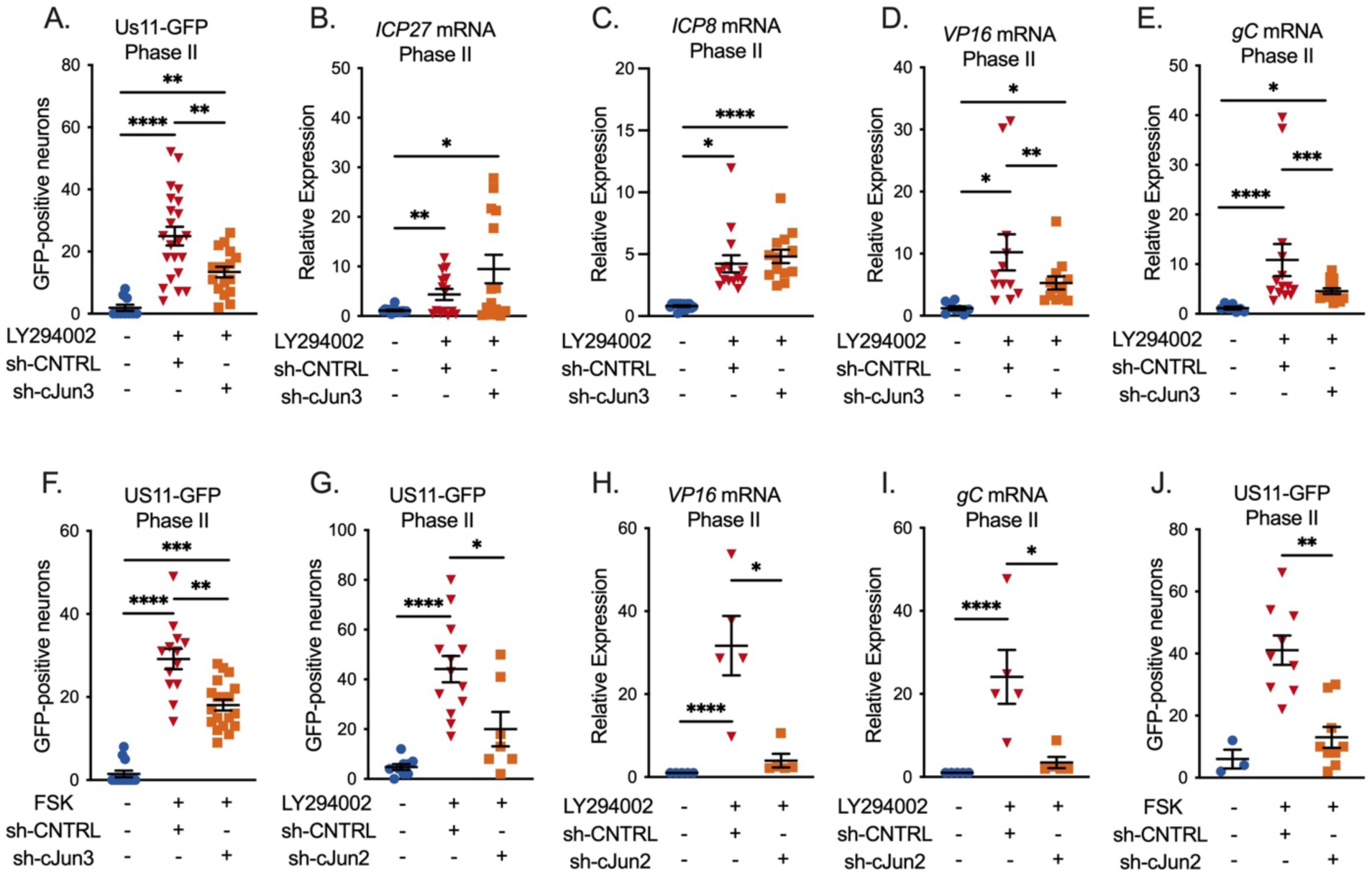
c-Jun is necessary for full HSV-1 reactivation. (A-F) Neurons were infected with HSV-1 and transduced at 6 days post-infection with a non-targeting shRNA lentivirus or sh-cJun3 (A-F) or sh-cJun2 (G-J) and reactivated 5 days later. Acyclovir was added for the first six days post-infection. (A,G) Quantification of Us11-GFP-positive neurons following reactivation with LY294002. (B-E, H-I) RT-qPCR for viral mRNA transcripts ICP27 (B), ICP8 (C), VP16 (D,H) gC (E,I), at 48 hours post-reactivation with LY294002. (F,J) Quantification of Us11-GFP-positive neurons following reactivation with forskolin. Individual replicates from at least 4 separate dissections are shown. Statistics determined by normal or non-normal (Wilcoxon-test, A, F-J) paired T-test. Individual biological replicates along with the means and SEMs are represented. * P < 0.05; ** P < 0.01. ns, not significant.

### c-Jun is required for full lytic replication in neurons

Phase II reactivation has previously been demonstrated to transcriptionally mirror HSV-1 *de novo* lytic infection in non-neuronal and neuronal cells, which contrasts with Phase I. Therefore, we investigated the contribution of c-Jun to *de novo* lytic infection in sympathetic neurons. Following c-Jun depletion and infection with HSV-1 Stayput-GFP, we quantified the numbers of Us11-GFP-positive neurons at 48 hours post-infection. This timepoint was chosen as it is when we previously detected the maximum number of GFP-positive neurons following *de novo* infection with Stayput-GFP (21). Consistent with our observation that c-Jun is required for full HSV-1 reactivation, the number of GFP-positive neurons indicative of *de novo* lytic infection events in this model system was robustly reduced following infection in c-Jun-depleted cultures (Fig 4A). In addition, viral DNA replication (Fig 4B) and transcription of all three classes of lytic genes, IE ICP27, E ICP8, L VP16, and TL gC and gM were robustly decreased in c-Jun depleted cells (Fig 4C), demonstrating that c-Jun is required for *de novo* lytic infection in neurons. These data suggest that c-Jun is a critical mediator of HSV-1 reactivation and lytic infection in neurons. However, these data also indicate that the mechanism of action may differ in reactivation versus *de novo* infection as c-Jun was required for expression of all three classes of viral genes during *de novo* infection but not during reactivation.

**Figure 4:**
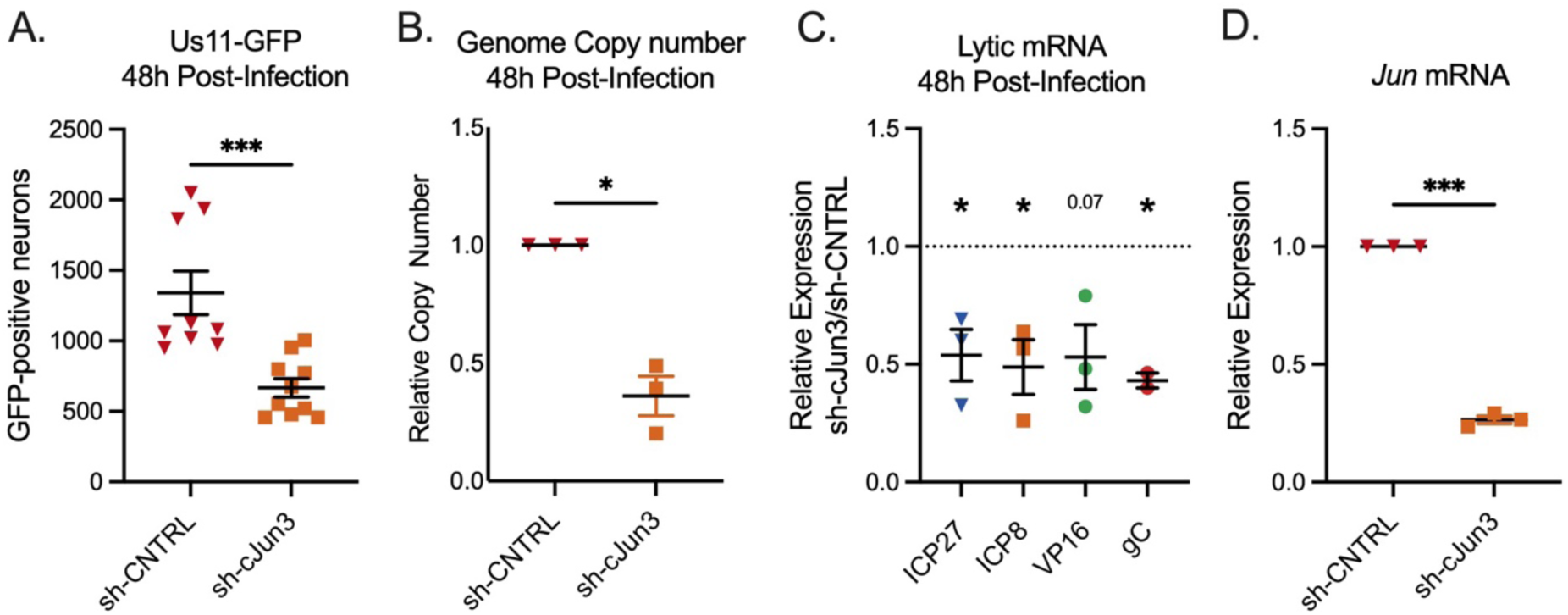
c-Jun is necessary for maximum *de novo* lytic infection in neurons. (A-H) Neurons were transduced with a non-targeting shRNA lentivirus or sh-cJun3 and infected with HSV-1 Stayput-GFP in the absence of viral DNA replication inhibitors 5 days post-transduction at an MOI of 5 PFU/cell for 48 hours. (A) Quantification of the numbers of GFP-positive neurons. Replicates from 3 separate dissections are shown; Unpaired t-test. (B) qPCR for viral DNA copy number. (C) RT-qPCR for viral mRNAs ICP27, ICP8, VP16, and gC. (D) RT-qPCR for Jun. Replicates from 3 separate dissections are shown; Paired t-test. Individual biological replicates along with the means and SEMs are represented. * P < 0.05; ** P < 0.01.

### c-Jun is present in HSV-1 replication compartments during full reactivation

As a DNA-binding protein, we proposed that c-Jun could modify viral gene expression during reactivation either directly on the viral genome or indirectly by altering a host cell factor. We therefore employed a single-cell imaging approach where the co-localization of c-Jun with individual reactivating neurons could be analyzed. As anticipated, c-Jun was not co-localized with latent viral genomes prior to the addition of the reactivation stimulus LY294002 (data not shown). To quantify c-Jun co-localization with viral genomes during Phase II reactivation, neurons were pulsed with 5-Ethynyl-2’-deoxycytidine (EdC; 10 μM) for 1 hour prior to carrying out Click-Chemistry to visualize viral genomes, along with immunofluorescence for c-Jun. We observed a robust up-regulation of c-Jun only in reactivating neurons (Fig 5A). Importantly, c-Jun robustly co-localized with replicating viral genomes during Phase II of reactivation (Fig 5A). This co-localization was quantified using a Pearson’s coefficient between c-Jun and EdC-Pulse (Fig 5B). Therefore, c-Jun is both up-regulated specifically in reactivating neurons and is recruited into viral replication compartments.

**Figure 5:**
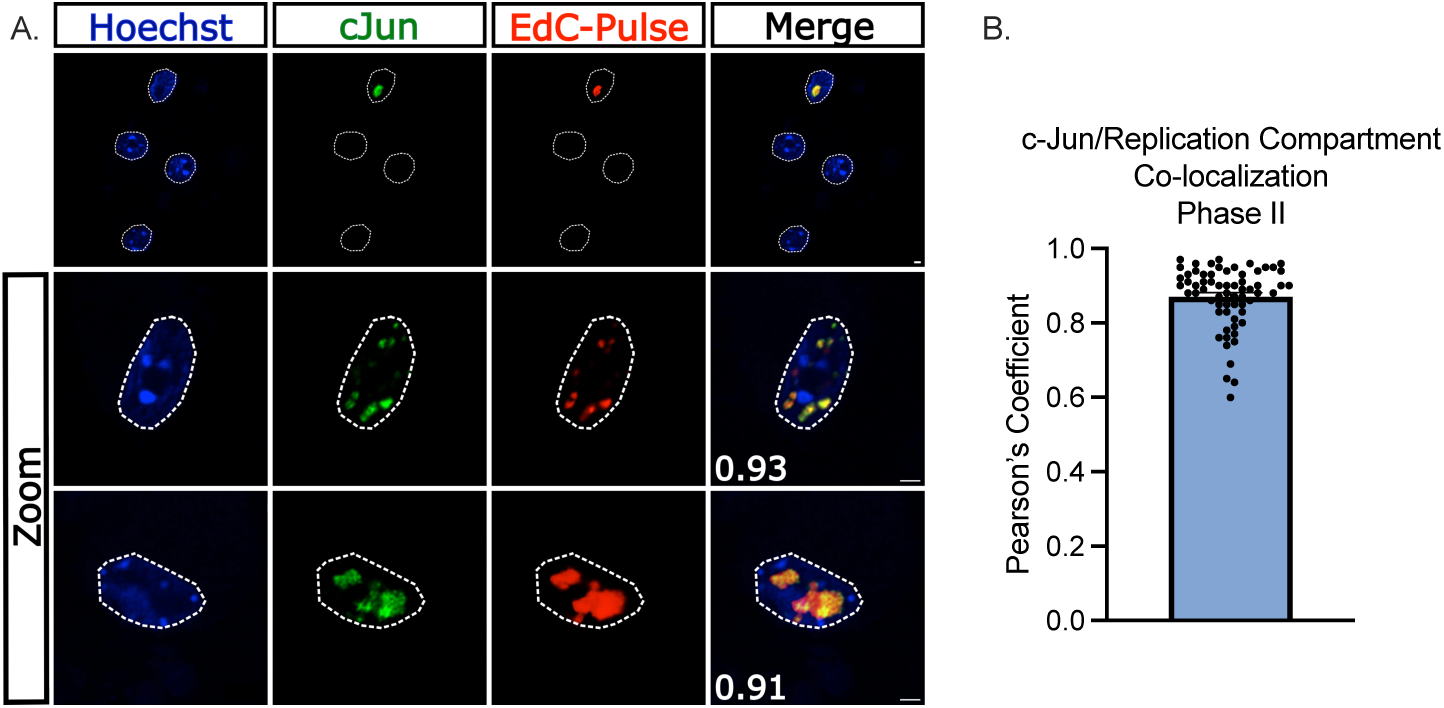
c-Jun colocalizes with replicating viral DNA during HSV-1 reactivation. (A-B) Latently infected neurons were reactivated with LY294002 and pulsed with 10 μM EdC for 1 hour to label viral DNA replication compartments. Nuclear stain Hoechst is shown in blue, and immunofluorescence was performed to visualize c-Jun in green. EdC-Pulse was visualized using click chemistry (shown in red). (A) Representative images of reactivating neurons. Scale bar = 10 μm. Pearson’s coefficient between c-Jun and EdC-Pulse featured in bottom left corner of merge. (B) Compiled Pearson’s coefficients. 35 individual neurons; 2 biological replicates.

### Activation of neuronal stress signaling during latency establishment enhances future reactivation

So far, our data point to a role for c-Jun specifically in stimulating late gene expression during Phase II reactivation. However, an additional observation was the differential phenotypes when c-Jun was depleted before infection versus before reactivation. This differential impact suggested that c-Jun signaling during initial infection promoted a form of viral latency that was more amenable to reactivation. As c-Jun is activated in response to neuronal stress signaling, we investigated how the active manipulation cell stress pathways and therefore enhanced c-Jun activation during initial infection impacted the later ability of the virus to reactivate. Inoculum with or without HSV-1 Stayput-Us11-GFP was added to neonatal sympathetic neurons in the presence or absence of nerve growth factor (NGF) for 3.5 hours. Immediately following the removal of this inoculum, the neurons were fixed and assayed for the presence of nuclear-localized and phosphorylated c-Jun (ser63), as an indication of c-Jun activation. The percentage of neurons under each condition demonstrating nuclear localized phosphorylated c-Jun was quantified from multiple fields of view, along with the mean intensity of the nuclear staining of each neuron. Using this approach, we were able to verify that NGF deprivation during the inoculation period is sufficient to elicit c-Jun activation, as anticipated (Fig. 6A-B). Interestingly, the neurons in the cultures infected with HSV-1 had enhanced c-Jun phosphorylation in both NGF-positive and -negative conditions as measured by the percentage of p-c-Jun-positive (Fig 6C). Therefore, HSV-1 *de novo* infection can also promote neuronal stress signaling indicated by c-Jun phosphorylation, which was also consistent with c-Jun activation during Phase II reactivation.

**Figure 6:**
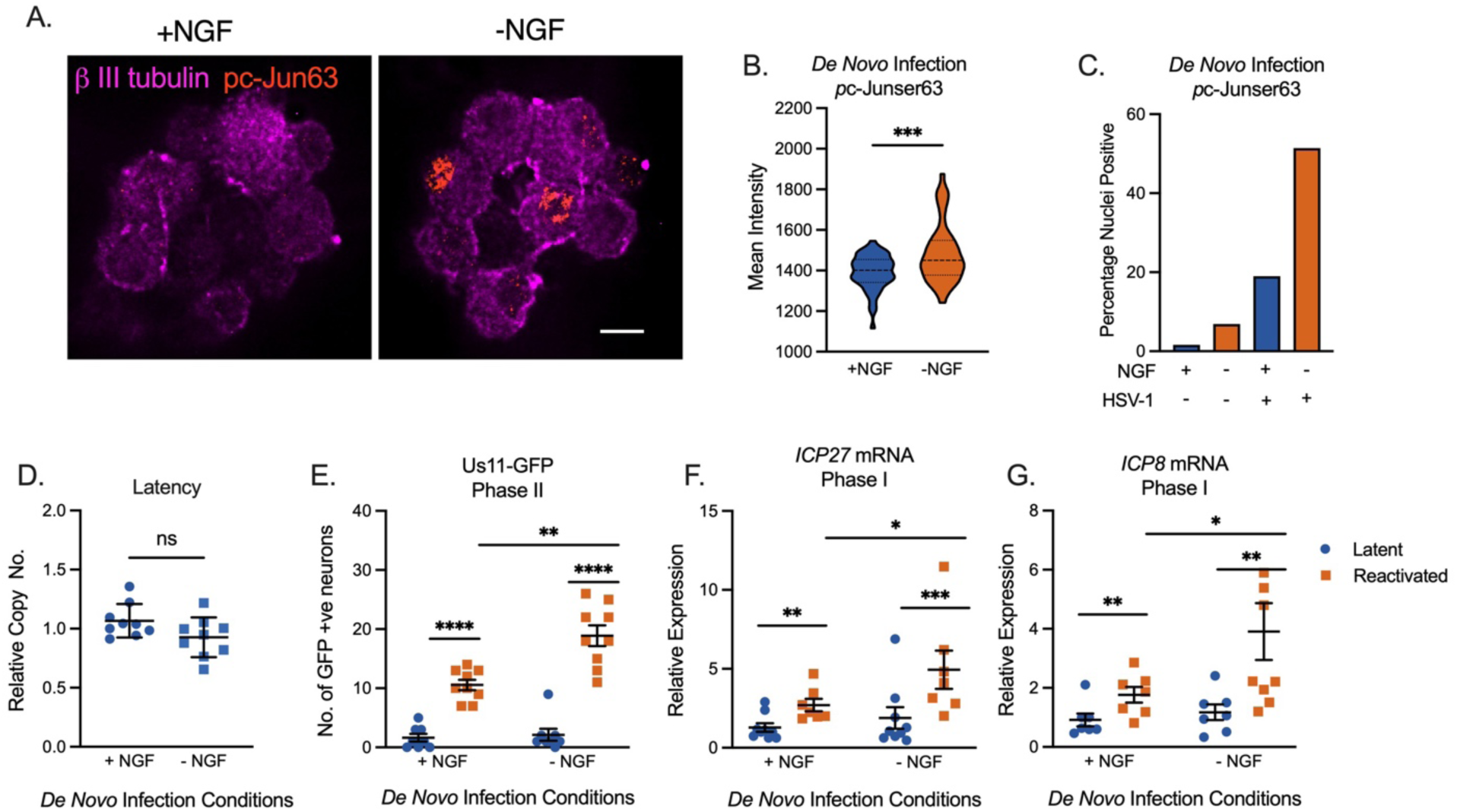
Stress Signaling Events during De Novo HSV-1 Infection. (A-C) Neonatal sympathetic cultures were infected with Stayput-GFP in inoculation media with or without nerve growth factor (NGF) for 3.5 hours and subsequently fixed and stained for neuronal marker B III tubulin (magenta) or phosphorylated c-Jun (orange). Representative image of nuclear phosphorylated c-Jun is demonstrated Scale bar = 10 μm. (B) Mean signal intensity for phosphorylated c-Jun in the nucleus following infection. N=100 from 1 biological replicate. (C) Quantification of proportion of neurons with pc-Jun-positive nuclei pooled from several fields of view. (D-G) Neuronal cultures were latently infected. NGF was either included or omitted during the 3.5 hours inoculation period. Cultures were later reactivated with LY294002 20 μM. (D) Latent viral DNA load. Replicates from 3 dissections shown. The peak number of GFP-positive neurons 48 hours post-stimulus (E) and relative expression of ICP27 (F) or ICP8 (G) transcripts 18 hours post-stimulus were quantified to analyze full reactivation and Phase I gene expression, respectively. Replicates from 3 dissections shown. Statistical comparisons were made using normal or non-normal (Mann-Whitney, E) t-Test. Individual biological replicates along with the means and SEMs are represented. * P < 0.05; ** P < 0.01.

Following our validation that c-Jun phosphorylation could be enhanced during initial infection by omitting NGF from the inoculum, we investigated the impact on HSV-1 reactivation. In agreement with the pre-latency establishment c-Jun depletion experiments (Fig 1), the perturbation of the NGF signaling pathway during initial infection did not alter latent viral DNA load (Fig 6D). However, in cultures where NGF had been deprived during the inoculation period and where c-Jun activation was enhanced, we observed enhanced reactivation stimulated with LY294002 based on the quantification of Us11-GFP-positive neurons at 48 hours post-treatment (Fig 6E). Importantly, Phase I gene expression, as analyzed through ICP27 and ICP8 expression 18 hours post-stimulus, was also enhanced in the cultures infected under NGF-deprivation conditions (Fig 6F-G). Therefore, these data indicate that enhanced neuronal stress mediated by reduced NGF-signaling and enhanced c-Jun phosphorylation, results in an enhanced ability of HSV-1 to ultimately undergo reactivation without impacting the latent viral load.

## Discussion

c-Jun is both a transcription and pioneer factor that is known to regulate host gene expression in neurons following cell stress. The up-regulation and phosphorylation of c-Jun following DLK-mediated activation of JNK is a key step in both neuronal apoptosis and axon pruning following the interruption of nerve growth factor signaling (50, 57, 58). The up-regulation of c-Jun-dependent genes is consistent with the kinetics of HSV-1 Phase I gene expression (59), hence our initial hypothesis that c-Jun plays a direct role in HSV-1 Phase I gene expression. However, our data indicate that c-Jun functions to promote reactivation but not during the JNK-dependent Phase I wave of lytic gene expression. In carrying out this study, we also observed that cell stress conditions during *de novo* infection have a long-term impact on either neurons themselves or the viral genome resulting in an enhanced ability of the virus to reactivate. This has important implications for how neurons have a memory of previous cellular stresses and for interpretations of HSV reactivation studies.

Although not required for Phase I gene expression, we did identify a role for c-Jun in promoting late gene expression during full, Phase II reactivation. Notably, this finding was also distinct from the role of c-Jun in promoting the expression of all three classes of viral genes during *de novo* infection. The exact mechanism of action and explanation for this difference between *de novo* infection and reactivation are unclear. One possible difference is the nature of the viral chromatin during reactivation versus *de novo* infection. The starting point for reactivation is a genome with a regularly spaced nucleosomal structure (60) and associated with heterochromatin (13, 18), whereas the viral genome during *de novo* infection is highly accessible and lacks a regular nucleosomal structure (1, 35, 61–64). Although the nature of the nucleosomal structure and overall accessibility of the viral genome during Phase II is currently unknown, it is possible that prior to viral DNA replication potential c-Jun binding sites are inaccessible. Additional proteins may also be activated in response to reactivation triggers that act during Phase I and on IE and E genes during Phase II as discussed below.

Because we were unable to detect a role for c-Jun during Phase I gene expression and IE/E gene expression during Phase II, this suggests additional undetermined host factors instead play a role. Additional candidates known to be activated or induced in response to reactivation triggers include other bZIP proteins such as Fos, JunD, ATF3 and Ddit3 (CHOP), along with other transcription factors; NF-Y, Gadd45α, Gadd45γ, and FOXO (59). In a previous study, depletion of Gadd45α had no impact on HSV-1 reactivation (65). An intriguing candidate for promoting Phase I gene expression is NF-Y complex because it has been implicated in the recruitment of JNK to chromatin during neuronal differentiation (66), and NF-Y has been identified as potentially stimulating transcription of the IE gene ICP0 in response to heat-stress (67), making NF-Y a viable candidate for driving initial reactivation events. Additional pioneer factors that may have a role in stimulating viral gene expression during reactivation are the Krüppel-like transcription factors (KLF) proteins, particularly KLF15 and KLF4 (68). KLF family members are up-regulated in corticosteroid-mediated reactivation of the related Bovine Herpesvirus-1 and may transactive viral immediate-early promoters (69–71). KLF4 is a well-known pioneer factor that activates previously silenced genes, a role that has been well characterized during cellular reprogramming (72, 73). The potential role of these proteins in regulating the transcription of HSV-1 genes from silenced heterochromatin during the different stages of HSV-1 reactivation therefore warrants further investigation.

c-Jun functions through either homo- or heterodimerization. Whether c-Jun binds to an additional bZIP protein to promote late gene expression is unknown. c-Jun can dimerize with Fos, Jun, CREB, or ATF family members. Recently, we found that JNK/DLK-dependent reactivation mediated by forskolin required CREB, as the addition of CREB inhibitor 666-15 restricted full reactivation as quantified by the number of GFP-positive neurons (28), although these data come with the caveat that a role for CREB has not been validated using genetic approaches. As observed here for c-Jun, CREB activity was not required for Phase I gene expression. Further, mapping the exact binding sites on the viral genome will help identify underlying sequence motifs bound by c-Jun, which can vary depending on the interacting protein (74). We did attempt to perform Cleavage Under Target & Release Under Nuclease (CUT&RUN) for c-Jun during Phase II reactivation. However, we found that the background signal for viral genomes from the non-specific control antibody was much higher on Phase II reactivating neurons than during latent infection (data not shown). For reasons that are not clear, ongoing viral DNA replication may result in substantial background in the CUT&RUN reaction, and therefore resolving c-Jun interacting sites on replicating viral genomes is currently problematic.

We also report a role for c-Jun for maximal *de novo* lytic infection in neurons as indicated by decreased of all classes of viral lytic genes, viral DNA replication, and late viral protein synthesis. Consistent with previous reports investigating lytic replication in non-neuronal cells, we found that HSV-1 infection and reactivation induced c-Jun activation (75, 76). However, a direct role for c-Jun during lytic replication has not previously been reported. Interestingly, a previous study from our lab found that JNK inhibition during *de novo* infection in neurons did not impact immediate early viral gene expression, although viral replication and the expression of other later classes of viral genes were not explored in this study (25). There, it remains possible that during *de novo* infection, c-Jun may be activated through an alternative pathway.

Environmental stressors have long been reported to correlate with HSV-1 reactivation and clinical disease. Fewer studies have explored the impact on HSV-1 and clinical outcomes when stress occurs during initial inoculation. Evidence from mouse models suggest that psychological stress during inoculation with HSV-1 enhances acute infection, as measured by infectious titer and pathology (77, 78). Complementary evidence from primary autonomic neurons similarly demonstrates elevated acute viral DNA replication and infectious virus production following HSV-1 infection in combination with stress hormone epinephrine (79). However, the impacts of stress during initial infection on later reactivation until now have not yet been investigated. Our findings suggest that additional stress that enhances c-Jun signaling during initial infection could exacerbate future reactivation. This has important implications for understanding the contributions to clinical HSV disease and why certain individuals may be more prone to reactivation than others. In addition, this observation has important experimental implications because it means that any manipulations performed on the virus or host during latency establishment could have an indirect effect on reactivation, and ideally the contribution of viral and host factors should be studied solely during reactivation to draw meaningful conclusions on their direct effects.

How the viral genome or neuron itself retains a memory of the initial infection conditions remains unclear. We previously found that viral genomes could retain a memory of interferon signaling during initial infection to result in restricted reactivation, which was mediated by association with repressive Promyelocytic Leukemia Nuclear Bodies (PML-NBs) (56). We now extend this study and show that cell stress has a converse effect and can prime future reactivation events. Given c-Jun’s ability to bind DNA and navigate chromatinized environments, it is tempting to speculate that stress signaling through c-Jun modifies the epigenetic nature of viral and/or host genomes to potentially induce a form of silencing that is more primed for transcriptional activation. Outside of the context of viral infection, early life stress has been implicated in leaving such a “chromatin scar” in the central nervous system, with changes in epigenetic signatures particularly for H3K27 and H3K79 methylation (80, 81) and a dysregulated priming of genes. Further studies on the mechanism of HSV latent infection and changes in the host and viral epigenetics will be important to understand how both the virus, and potentially neurons themselves, have a memory of previous cell stress or immune events and the impact on future responses.

## Materials and Methods

### Preparation of HSV-1 virus stocks

Stocks of Stayput Us11-GFP (strain SC16) for *in vitro* experiments were propagated and titrated on gH-complementing F6 cells (21, 82). Vero F6 cells were maintained in Dulbecco’s modified Eagle’s medium (Gibco) supplemented with 10% FetalPlex (Gemini Bio-Products) and 250 μg/mL of G418/Geneticin (Gibco).

### Primary neuronal cultures

Sympathetic neurons from the Superior Cervical Ganglia (SCG) of post-natal day 0-2 (P0-P2) CD1 Mice (Charles River Laboratories) were dissected as previously described (25). Rodent handling and husbandry were carried out under animal protocols approved by the Animal Care and Use Committee of the University of Virginia (UVA). Ganglia were briefly kept in Leibovitz’s L-15 media with 2.05 mM l-glutamine before dissociation in collagenase type IV (1 mg/ml) followed by trypsin (2.5 mg/ml) for 20 min; each dissociation step was at 37°C. Dissociated ganglia were triturated, and approximately 10,000 neurons per well were plated onto rat tail collagen in a 24-well plate. Sympathetic neurons were maintained in feeding media: Neurobasal® Medium supplemented with PRIME-XV IS21 Neuronal Supplement (Irvine Scientific), 50 ng/ml Mouse NGF 2.5S (Alomone labs), 2 mM l-Glutamine, and 100 µg/ml Primocin (Invivogen). Aphidicolin (3.3 µg/ml) was added to the media for the first five days post-dissection to select against proliferating cells.

### Lytic HSV-1 infection in primary neurons

P6-8 SCG neurons were infected with Stayput Us11-GFP at MOI 5 PFU/cell, (assuming 10,000 cells per well) in Dulbecco’s Phosphate Buffered Saline (DPBS) + CaCl2 + MgCl2 supplemented with 1% fetal bovine serum and 4.5 g/L glucose for 3.5 h at 37°C. The inoculum was replaced with feeding media (as described above). Acyclovir (ACV) was not utilized in these infections. Lytic infection was quantified by quantifying the numbers of GFP-positive neurons.

### Establishment and reactivation of latent HSV-1 infection in primary neurons

P6-8 SCG neurons were infected with Stayput Us11-GFP at MOI 7.5 PFU/cell, (assuming 10,000 cells per well) in Dulbecco’s Phosphate Buffered Saline (DPBS) + CaCl2 + MgCl2 supplemented with 1% fetal bovine serum, 4.5 g/L glucose, and 10 μM acyclovir (ACV) for 3.5 h at 37°C. The inoculum was replaced with feeding media (as described above) with 50 μM ACV. 6 days post-infection, ACV was washed out and replaced with feeding media alone. Reactivation was reported by quantifying the numbers of GFP-positive neurons following the addition of 20-40 µM LY294002 (Tocris) or 60 µM forskolin (Tocris).

### Analysis of viral DNA load and mRNA expression by reverse transcription– quantitative PCR (RT–qPCR)

To assess the relative expression of HSV-1 lytic mRNA, total RNA was extracted from approximately 10,000 neurons using the Quick-RNA™ Miniprep Kit (Zymo Research) with an on-column DNase I digestion. mRNA was converted to cDNA using the Maxima First Strand cDNA Synthesis Kit for RT-qPCR (Fisher Scientific), using random hexamers for first-strand synthesis and equal amounts of RNA (20–30 ng/reaction). To assess viral DNA load, total DNA was extracted from approximately 10,000 neurons using the Quick-DNA™ Miniprep Plus Kit (Zymo Research). qPCR was carried out using PowerUp™ SYBR™ Green Master Mix (ThermoFish Scientific). The relative mRNA or DNA copy number was determined using the comparative CT (ΔΔCT) method normalized to mRNA or DNA levels in latently infected samples. Viral RNAs were normalized to mouse reference gene mGAPDH RNA. All samples were run in triplicate on an Applied Biosystems™ QuantStudio™ 6 Flex Real-Time PCR System and the mean fold change compared to the calculated reference gene. The sequence of both forward and reverse primers used have been published previously (28).

### Preparation of lentiviral vectors

Lentiviruses expressing shRNA against c-Jun (c-Jun-1 = TRCN0000360511, c-Jun-2 = TRCN0000055205, c-Jun-3 = TRCN0000042696) or a control lentivirus shRNA (pLKO.1 vector expressing a non-targeting shRNA control) were prepared by co-transfection using JetPRIME® with psPAX2 and pCMV-VSV-G (83) into the 293LTV packaging cell line (Cell Biolabs). Supernatant was harvested at 40- and 64-h post-transfection and filtered using a 45 μM PES filter. Sympathetic neurons were transduced overnight in neuronal media containing 8 μg/ml protamine sulfate and 50 μM ACV.

### Western Blotting

Neurons were lysed in RIPA Buffer with cOmplete, Mini, EDTA-Free Protease Inhibitor Cocktail (Roche) and PhosSTOP Phosphatase Inhibitor Cocktail (Roche) on ice for 2 hours with regular vortexing to aid lysis. Insoluble proteins were removed via centrifugation, and lysate protein concentration was determined using the Pierce Bicinchoninic Acid Protein Assay Kit (Invitrogen) using a standard curve created with BSA standards of known concentration. Equal quantities of protein (15–50 μg) were resolved on 4–20% gradient SDS-Polyacrylamide gels (Bio-Rad) and then transferred onto Polyvinylidene difluoride membranes (Millipore Sigma). Membranes were blocked in PVDF Blocking Reagent for Can Get Signal (Toyobo) for 1 hr. Primary antibodies were diluted in Can Get Signal Immunoreaction Enhancer Solution 1 (Toyobo) and membranes were incubated overnight at 4°C. HRP-labeled secondary antibodies were diluted in Can Get Signal Immunoreaction Enhancer Solution 2 (Toyobo) and membranes were incubated for 1 hr at room temperature. Antibody usage is recorded in Table 1. Blots were developed using Western Lightning Plus-ECL Enhanced Chemiluminescence Substrate (PerkinElmer) and ProSignal ECL Blotting Film (Prometheus Protein Biology Products) according to manufacturer’s instructions. Blots were stripped for reblotting using NewBlot PVDF Stripping Buffer (Licor). Band density was quantified in ImageJ.

**Table 1.** Antibodies.

| **Table 1. Antibodies** |  |  |  |
| --- | --- | --- | --- |
| **Designation** | **Source or reference** | **Identifiers** | **Additional Information** |
| Anti-c-Jun (Mouse monoclonal) | BD Biosciences | Cat #610326 | WB (1:1,000) |
| Anti-α-tubulin (Mouse monoclonal) | Millipore sigma | Cat #T9026 | WB (1:2500) |
| Anti-Rabbit IgG Antibody (H+L), Peroxidase (Goat polyclonal) | Vector Labs | Cat #PI-1000 | WB (1:10,000) |
| Anti-mouse IgG Antibody (H+L), Peroxidase (Horse polyclonal) | Vector Labs | Cat #PI-2000 | WB (1:10,000) |
| Phospho-c-Jun (Ser63) II (Rabbit polyclonal) | Cell Signaling Technology | Cat #9261 | IF (1: 400) |
| Beta III Tubulin (Chicken polyclonal) | EMD Millipore | Cat #9354 | IF (1:500) |

### Immunofluorescence

Neurons were fixed for 15 min in 4% formaldehyde and blocked for 1 hour in 5% bovine serum albumin and 0.3% Triton X-100, and incubated overnight in primary antibody. Following primary antibody treatment, neurons were incubated for 1 h in Alexa Fluor 488-, 555-, and 647-conjugated secondary antibodies for multicolor imaging (Invitrogen). Nuclei were stained with Hoechst 33258 (Life Technologies). Images were acquired using an sCMOS charge-coupled device camera (pco.edge) mounted on a Nikon Eclipse Ti inverted epifluorescent microscope using NIS-Elements software (Nikon). Images were analyzed using ImageJ.

### Click Chemistry

To label replicating HSV-1 DNA, reactivated neuronal cultures were pulsed with 10 μM EdC for 1 hour prior to fixation and processing. Click-chemistry was carried out a described previously (84) with some modifications to visualize EdC-incorporated replicating viral genomes. Neurons were washed with CSK buffer (10 mM HEPES, 100 mM NaCl, 300 mM Sucrose, 3 mM MgCl2, 5 mM EGTA) and simultaneously fixed and permeabilized for 10 min in 1.8% methanol-free formaldehyde (0.5% Triton X-100, 1% phenylmethylsulfonyl fluoride (PMSF)) in CSK buffer, then washed twice with PBS before continuing to the click-chemistry reaction and immunostaining. Samples were blocked with 3% BSA for 30 min, followed by click-chemistry using EdC-labeled HSV-1 DNA and the Click-iT EdU Alexa Flour 555 Imaging Kit (ThermoFisher Scientific, C10638) according to the manufacturer’s instructions. For immunostaining, samples were incubated overnight with primary antibodies in 3% BSA. Following primary antibody treatment, neurons were incubated for 1 hr in Alexa Fluor 488-, 555-, and 647-conjugated secondary antibodies for multi-color imaging (Invitrogen). Nuclei were stained with Hoechst 33258 (Life Technologies). Images were acquired at 60x using an sCMOS charge-coupled device camera (pco.edge) mounted on a Nikon Eclipse Ti Inverted Epifluorescent microscope using NIS-Elements software (Nikon). Images were analyzed and intensity quantified using ImageJ.

## Statistical Analysis

Power analysis was used to determine the appropriate sample sizes for statistical analysis. All statistical analysis was performed using Prism V10. Normality of the data was determined with the Kolmogorov-Smirnov test. Specific analyses are included in the figure legends.

## Acknowledgments

We thank Gary Cohen at the University of Pennsylvania for Vero F6 cells. This work was supported by National Institutes of Health grants NS105630 (ARC), T32GM008136 (SAD and AKF), and T32AI007046 and The Owens Family Foundation (ARC).

## Notes

### Competing Interest Statement

The authors have declared no competing interest.

